# Sawfish: Improving long-read structural variant discovery and genotyping with local haplotype modeling

**DOI:** 10.1101/2024.08.19.608674

**Authors:** Christopher T. Saunders, James M. Holt, Daniel N. Baker, Juniper A. Lake, Jonathan R. Belyeu, Zev Kronenberg, William J. Rowell, Michael A. Eberle

## Abstract

**Motivation:** Structural variants (SVs) play an important role in evolutionary and functional genomics but are challenging to characterize. High-accuracy, long-read sequencing can substantially improve SV characterization when coupled with effective calling methods. While state-of the-art long-read SV callers are highly accurate, further improvements are achievable by systematically modeling local haplotypes during SV discovery and genotyping.

**Results:** We describe sawfish, an SV caller for mapped high-quality long reads incorporating systematic SV haplotype modeling to improve accuracy and resolution. Assessment against the draft Genome in a Bottle (GIAB) SV benchmark from the T2T-HG002-Q100 diploid assembly shows that sawfish has the highest accuracy among state-of-the-art long-read SV callers across every tested SV size group. Additionally, sawfish maintains the highest accuracy at every tested depth level from 10 to 32-fold coverage, such that other callers required at least 30-fold coverage to match sawfish accuracy at 15-fold coverage. Sawfish also shows the highest accuracy in the GIAB challenging medically relevant genes benchmark, demonstrating improvements in both comprehensive and medically relevant contexts.

When joint-genotyping 10 samples from CEPH-1463, sawfish has over 9000 more pedigree-concordant calls than other state-of-the-art SV callers, with the highest proportion of concordant SVs (78%) as well. Sawfish’s quality model can be used to select for an even higher proportion of concordant SVs (86%), while still calling over 5000 more pedigree-concordant SVs than other callers. These results demonstrate that sawfish improves on the state-of-the-art for long-read SV calling accuracy across both individual and joint-sample analyses.

**Availability:** Sawfish is released as a pre-compiled Linux binary and user guide on GitHub: https://github.com/PacificBiosciences/sawfish.

## Introduction

Structural variants (SVs) are medium to large-scale rearrangements of the genome, typically characterized as events at least 50 bases in size. Though there are relatively few SVs in the human genome compared to the number of single nucleotide variants and small indels, collectively these SVs impact more bases than small variants and play a significant role in human evolution and disease^1^. Unfortunately, SVs are often mediated by repetitive sequence that can be difficult to study with short-read sequencing technologies^2–4^, so our understanding of these variants has been limited compared to smaller variants. Recent advances in highly accurate long-read sequencing together with updated SV detection methods have greatly improved the quality of SVs routinely characterized from human samples^3,5^, but continued improvements to SV characterization methods are critical to realizing the full potential of long-read genome sequencing.

We describe a new method, sawfish, which assembles SV haplotypes and models variants at the haplotype level in all downstream sample merging and genotyping steps. We show that such an approach can substantially improve SV accuracy on the latest high-resolution assembly-based SV benchmarks. Compared to other state-of-the-art long-read SV callers on these benchmarks, sawfish provides the most accurate SV calls in aggregate as well as across SV size ranges. We also show that sawfish provides the most accurate SV calls at every tested sequencing depth from 10 to 32-fold coverage, allowing sawfish with 15-fold sequencing coverage to match the accuracy of other callers at 30-fold coverage. We additionally show that sawfish’s approach leads to higher genotype concordance on a large pedigree, where sawfish not only produces thousands more concordant SVs than other callers, but also a greater proportion of concordant SVs, indicative of a more accurate genotyping model. In aggregate, our results show that sawfish’s methods comprehensively improve the accuracy of long-read SV calling in both single and joint-sample analysis scenarios.

## Methods

Sawfish is capable of calling and genotyping deletions, insertions, duplications, translocations and inversions from mapped high-accuracy long reads. The method is designed to discover breakpoint evidence from each sample, then merge and genotype variant calls across samples in a subsequent joint-genotyping step, using a process that emphasizes representation of each SV’s local haplotype sequence to improve accuracy. The primary sawfish SV calling steps are to: (1) identify clusters of SV-associated alignment signatures from mapped sequencing reads; (2) assemble/polish SV-associated reads from each cluster into one or more haplotype consensus sequences; (3) merge putative SV haplotypes across samples; (4) align haplotypes back to a reference sequence; (5) genotype each SV via assessment of relative read support for each local haplotype; and (6) further refine SV type classification based on the supporting depth signature for unbalanced SV candidates. Details for each of these SV calling operations are described in Supplementary Methods.

In addition to using local haplotype assembly, all sawfish SVs are modeled and genotyped at the breakpoint level, which allows all breakpoints to be consistently reported at basepair resolution, with any breakpoint insertion and microhomology sequence detected and described in the output. For multi-breakpoint events, such as inversions, sawfish will report both the event and its component breakpoints to retain this breakpoint detail level for all cases. Another feature derived from the local haplotype modeling in sawfish is that local SV phasing can be reported in the output wherever two variant haplotypes overlap in the same sample, helping to provide a more detailed description of complex variation regions.

Due to the high-resolution of sawfish’s SV calls, and the tendency for this to result in overlapping SV alleles in complex variation regions, we assess single-sample call accuracy on HG002 against a recent similarly high-resolution benchmark, the draft GIAB SV benchmark based on the T2T-HG002-Q100 diploid assembly. This benchmark requires the use of updated assessment workflows to harmonize query and benchmark SV representation in complex variation regions, so we test with two independent assessment tools (Truvari^6^ and hap-eval^7^) to check for consistent qualitative assessment results across both tools (see Supplementary Methods). Although this strategy introduces additional assessment complexity, compared to the previous mapping-based GIAB SV benchmark^8^, the newer assembly-based benchmark contains roughly 3 times as many confident SVs (30244 vs 9646), and spot-checking of trial assessment results labeled as false positives or false negatives from each benchmark set has shown a much higher level of assessment artifact when using the previous mapping-based benchmark.

## Results

The local haplotype assembly implemented in sawfish leads to improved SV calling accuracy in both single-sample and multi-sample joint-genotyping contexts. Here we assess SV performance for both contexts compared to two state-of-the-art long-read SV callers, Sniffles2^9^ and pbsv^10^.

### Single-sample accuracy

To benchmark SV performance, we take advantage of recent advances from the HG002 Q100 project^11^, which seeks to create a near perfect diploid genome for HG002 based on long-read assembly. This high-quality diploid genome assembly can be used to generate a comprehensive benchmark by aligning assembly contigs back to a standard reference genome. A draft SV benchmark using this approach has been created by GIAB^12^ which we use to assess single-sample SV caller accuracy. Compared to the previous GIAB SV benchmark^8^, this assembly-based benchmark contains roughly 3 times as many confident SVs (30244 vs 9646) to more comprehensively assess SVs throughout the genome.

All SV callers are first assessed on recent HiFi WGS data for HG002 at 32-fold coverage to characterize accuracy at typical full WGS depth. We assess accuracy against the GIAB T2T SV benchmark using Truvari^6^ (see Supplementary Methods), finding that the sawfish F1-score of 0.974 is over 1.3% higher than both Sniffles2 and pbsv, as shown in **Table S1**. Given the recent development of the T2T SV benchmark and its associated assessment methods, we check this result by replicating the assessment with an independent method, hap-eval^7^ (see Supplementary Methods) and find that the observed accuracy trend is consistent across assessment methods. The hap-eval sawfish F1-score is 0.974, which in this case is over 3.7% and 4.4% higher than Sniffles2 and pbsv, respectively. After this assessment consistency check, all further analysis is consolidated to use Truvari.

We next stratify the accuracy benchmark by SV length into 3 size ranges: 50 to 499 bases, 500 to 4999 bases, and 5000 bases or more (**Figure 1a, Table S2**). We observe that sawfish has the highest F1-score in every size range, with a modest F1-score increase of 0.5% for the smallest (50-499 bp) SVs compared to the next-best method, pbsv, but with a growing lead as SV size increases. For SVs of at least 5000 bases, sawfish’s F1-score is over 3.7% and 7.9% higher than Sniffles2 and pbsv, respectively. Thus, we observe that the higher accuracy of sawfish calls extends across SV size ranges. When the benchmark SVs are further separated into insertions and deletions, in addition to size ranges, we continue to see sawfish retain the highest F1-score among the tested SV callers in every SV type category (**Table S2**).

**Fig. 1.**
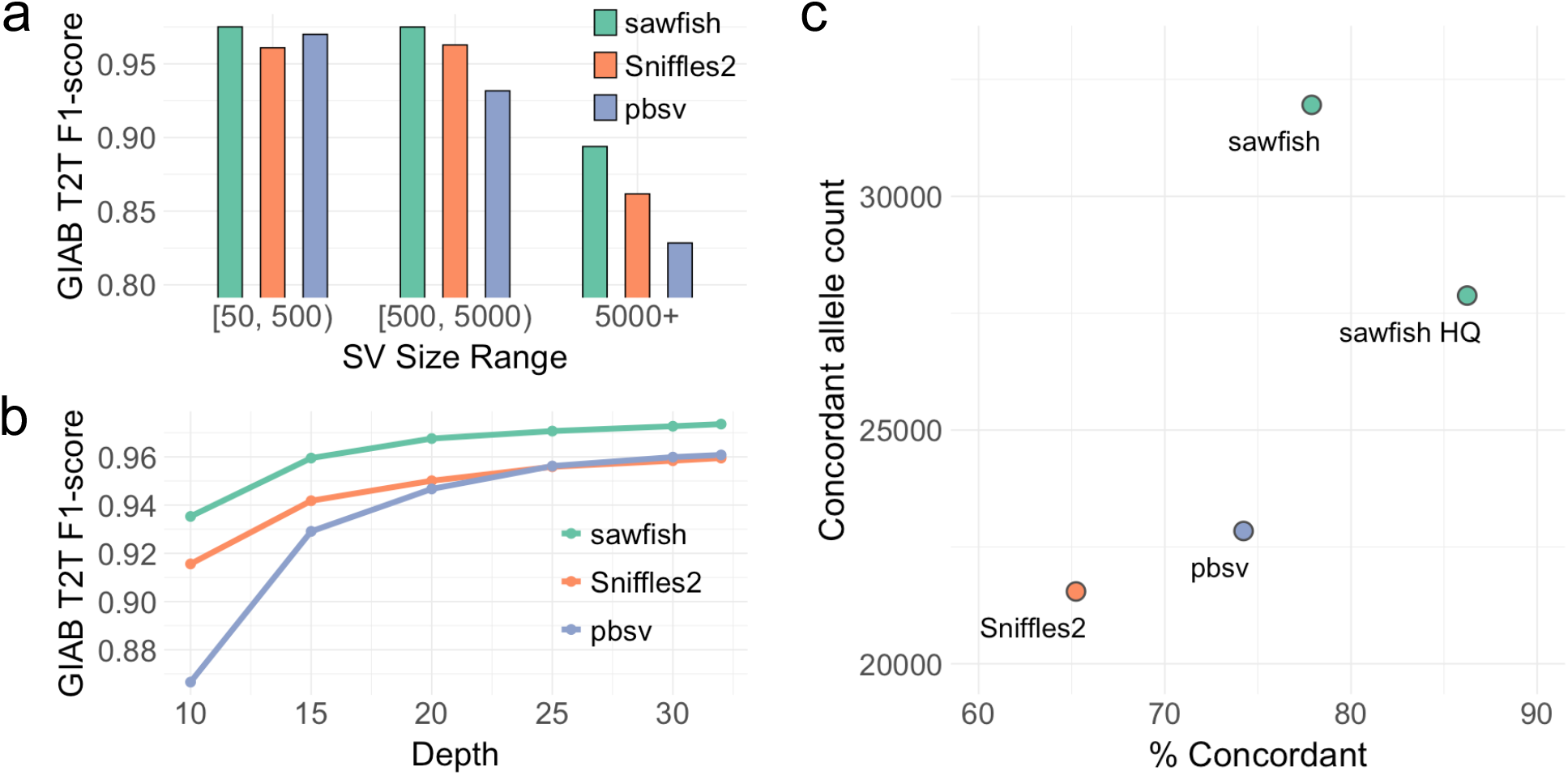
SV caller accuracy assessment. **a**. Assessment of SV caller performance on the HG002 GIAB draft T2T assembly-based benchmark for HiFi WGS data from HG002 at 32-fold coverage. Results are stratified by SV size, showing consistently improved F1-score for sawfish from smaller to larger SV sizes. **b**. HG002 SV caller performance assessed as in part (a) but for all SV sizes with depth levels subsampled down to 10-fold coverage, showing improved F1-score for sawfish at all coverage levels. **c**. Assessment of SV caller joint-genotyping on CEPH Pedigree 1463 generations 2 and 3 samples. For each SV caller the number of SVs where genotypes are concordant with the pedigree inheritance pattern across all samples are shown compared to the percentage of all SVs which are concordant. ‘sawfish HQ’ shows sawfish results filtered for genotype quality (GQ) >= 40 in all samples. The results show that sawfish calls thousands more concordant SVs than other SV callers while at the same time calling proportionally more concordant SVs, where the proportion of concordant SVs can be increased to over 86% with modest quality filtration that maintains a very high concordant SV count.

We additionally evaluate how read coverage influences SV call accuracy by running all SV callers on WGS data subsampled at various levels down to 10-fold coverage (**Figure 1b, Table S3**). We have already shown that sawfish has the highest accuracy at full coverage, and here we observe that as coverage is reduced sawfish outperforms other methods by even greater margins, at 10-fold coverage the sawfish F1-score is over 2.1% higher than Sniffles2 and 7.9% higher than pbsv. In fact, at 15-fold coverage sawfish achieves roughly the same accuracy as other SV callers at 30-fold coverage, suggesting that population studies could use lower depth without loss of SV accuracy through improved calling and genotyping methods.

### Medically relevant SV benchmark

We compare SV caller performance on the GIAB challenging medically relevant genes (CMRG) benchmark set of 217 SVs^13^. Although all long-read SV callers are relatively accurate in this context, we observe even higher accuracy for sawfish calls compared to other callers, with sawfish calls including only 2 false negatives and 1 false positive, resulting in an F1-score of 0.993 (**Table S4**). This compares to an F1-score of 0.964 for Sniffles and 0.971 for pbsv. This result demonstrates that sawfish methods improvements are not just applicable to comprehensive SV benchmark performance, but SVs in medically relevant genes as well.

## Genotype concordance in a large pedigree

While the sawfish haplotype modeling approach has been shown to benefit SV accuracy in individual samples, it also improves joint-genotyping accuracy across multiple samples via merging observations and genotyping at the SV haplotype level. By directly comparing SV haplotypes between samples in the merging process, the method reduces alignment-dependent artifacts introduced in the direct comparison of variants.

To assess joint-genotyping accuracy, we evaluate SVs from HiFi WGS data recently generated for all members of CEPH Pedigree 1463^14^. For each SV caller we jointly genotype SVs across 10 samples comprising the 2^nd^ and 3^rd^ pedigree generations. For each SV we can then evaluate whether the genotypes called across all samples are consistent with the known pedigree haplotype inheritance pattern in that region, as described in Eberle at al.^15^. We summarize the results of this analysis in **Figure 1c** and **Table S5**, which show that sawfish not only has 9115 more concordant SVs than either of the other SV callers, but it also has proportionally more concordant calls at over 77.8%.

To evaluate sawfish’s genotype quality score for this application, we filter sawfish calls for a genotype quality of at least 40 in all samples, creating a call set labeled “sawfish HQ” (**Figure 1c, Table S5**). The sawfish HQ call set still retains 5035 more concordant SVs than the next best caller, but the call set also has over 86.2% concordance, which is 12% greater than the next best method, pbsv. Sniffles2 also provides genotype quality values, so we filter the Sniffles2 calls to require a genotype quality of at least 10 for all samples to get a similar concordance (85.25%), but this requires filtering out almost 2/3 of the original Sniffles2 concordant SV call set, leaving only 7643 concordant SVs (**Table S5**). This result shows that sawfish’s quality model provides a more effective joint-genotyping solution to achieve both high sensitivity and genotype accuracy.

## Conclusion

Our assessment of long-read SV calling accuracy shows that sawfish’s approach to SV haplotype modeling consistently improves SV accuracy in both single-sample and multi-sample joint-genotyping contexts, and further demonstrates that this accuracy gain is consistent over SV size ranges and across typical WGS coverage levels. In a joint-genotyping context, sawfish calls many more concordant SVs than other callers, while providing a higher enrichment for concordance among all calls. In addition to the advantages of SV haplotype modeling, sawfish’s high accuracy shows the advantages of supporting depth evaluation for large deletions and duplications, native phasing for proximal or overlapping SVs, and an effective genotype quality model which allows for flexible filtration options. These methods will continue to be developed and refined against newly available high-quality assembly-based SV benchmark sets to help realize the full potential of SV discovery through comprehensive long-read genome sequencing.

## Acknowledgements

We would like to thank early sawfish users for their valuable feedback and suggestions, including detailed feedback from Wolfram Höps and Christian Gilissen leading to several methods improvements. We thank Nathanael Olson and Justin Zook for providing early access and help with the T2T-HG002-Q100 draft SV benchmark, as well as Adam English and Donald Freed for their assistance with using and interpreting SV assessment tools.

## Supplementary Material

### Supplementary Methods

#### Single sample SV accuracy assessment

##### Sequencing Data Processing

We used publicly available HiFi WGS data for HG002, sequenced to ∼32-fold coverage on the Revio system. The unmapped sequencing data can be downloaded from:

https://downloads.pacbcloud.com/public/revio/2022Q4/HG002-rep1/m84011_220902_175841_s1.hifi_reads.bam

The sequencing data is mapped to GRCh38 using pbmm2 version 1.13.1 with the ‘CCS’ preset option. The mapped reads were subsampled to each test coverage level shown in **Figure 1a** using the subsample option in samtools^16^ version 1.17 based on mapped coverage assessment from mosdepth^17^ version 0.2.6.

##### SV calling

The two SV callers, Sniffles2^9^ and pbsv^10^, used for comparison in this study were selected based on the following criteria for our single and multi-sample accuracy assessment: (1) support for SV-calling from long reads; (2) SV genotyping; (3) prediction of all common SV types, including deletions, insertions, duplications, breakpoints and inversions; and (4) direct facility to joint-genotype SVs over multiple samples.

SV calls were generated from sawfish version 0.12.0, Sniffles2 version 2.3.3, and pbsv version 2.9.0. The command-line templates for single-sample analysis of each SV caller are as follows:

###### Program 1 | Command line template for sawfish single-sample calling

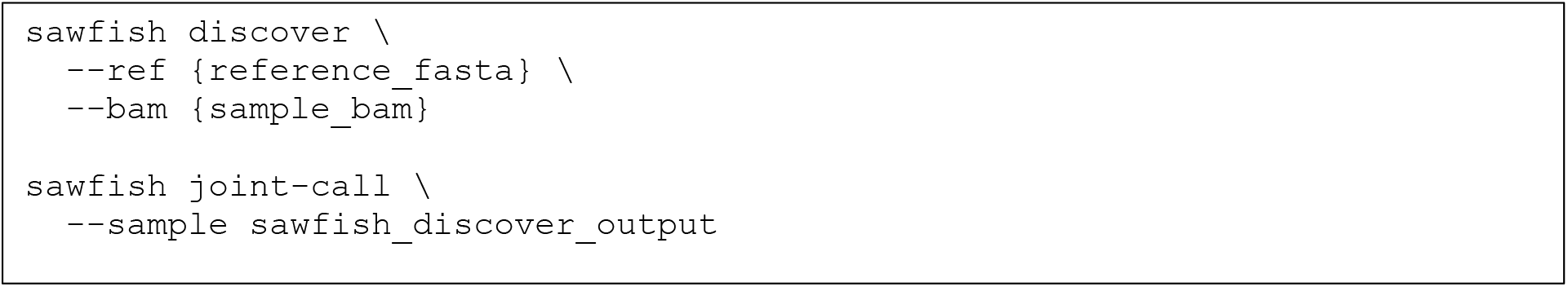

###### Program 2 | Command line template for Sniffles2 single-sample calling

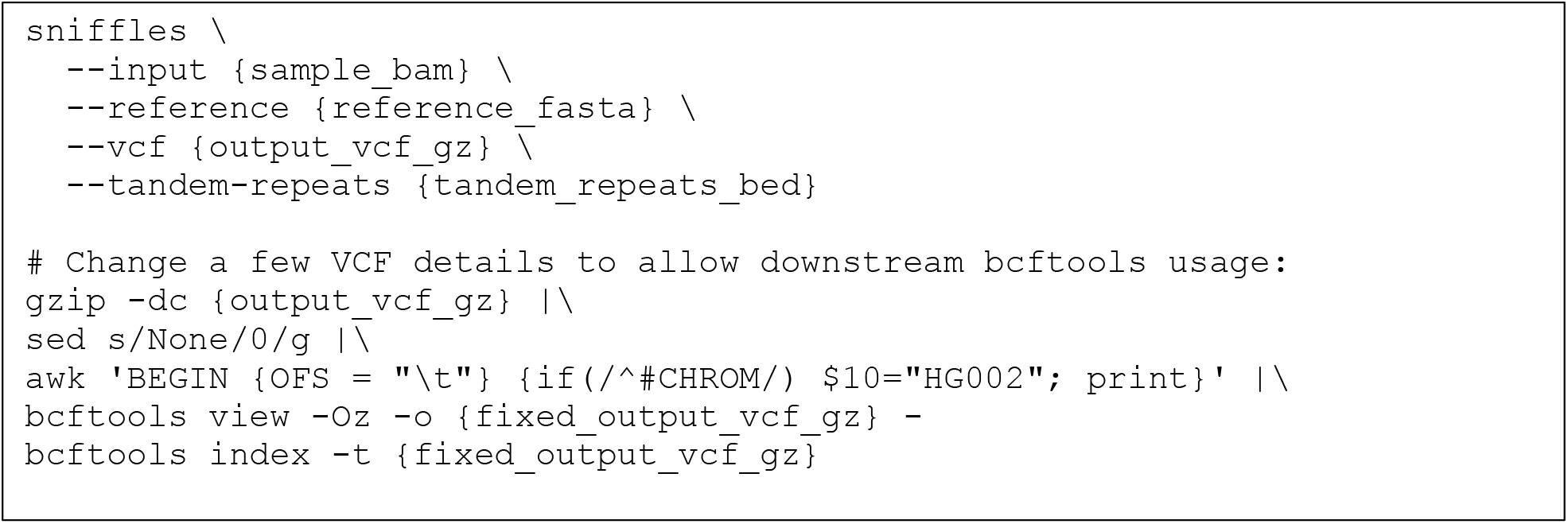

###### Program 3 | Command line template for pbsv single-sample calling

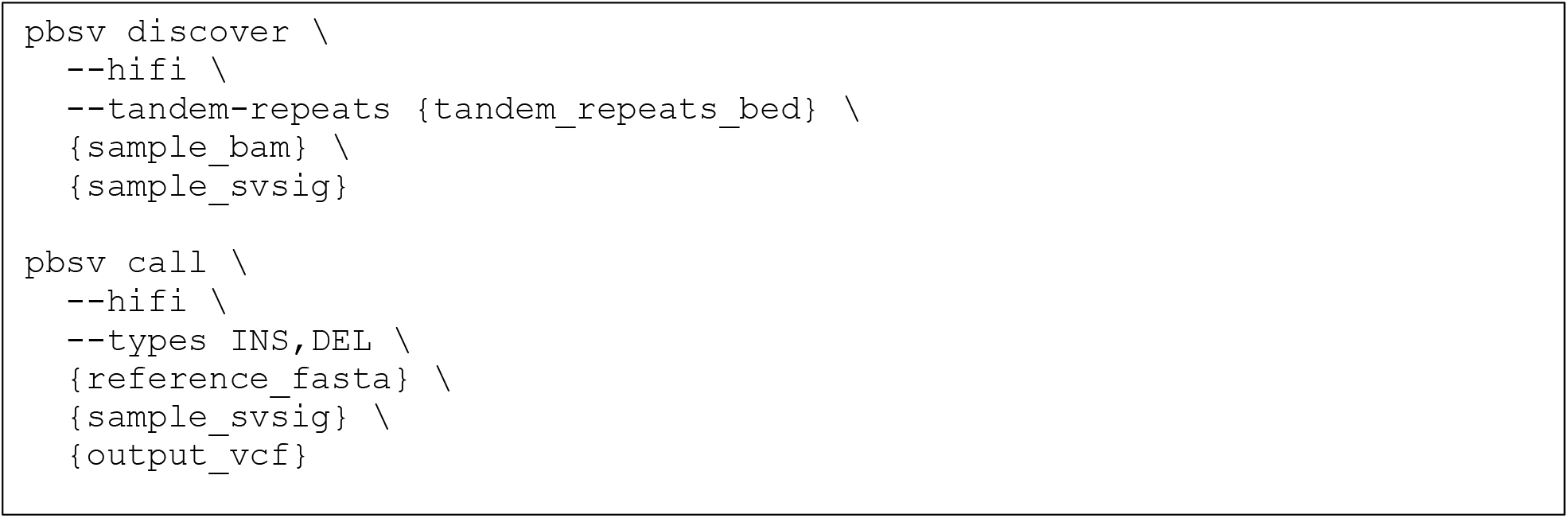

##### Assessment of SV calls against the GIAB draft SV benchmark

Comprehensive assessment of SV calls on HG002 was conducted against the GIAB draft SV benchmark (V0.015-20240215) based on the T2T-HG002-Q100v1.0 diploid assembly aligned to GRCh38. The benchmark variants and confidence regions were downloaded from:

https://ftp-trace.ncbi.nlm.nih.gov/ReferenceSamples/giab/data/AshkenazimTrio/analysis/NIST_HG002_DraftBenchmark_defrabbV0.015-20240215

This benchmark was modified to exclude sex chromosomes by removing chromosomes X and Y from the confidence regions and additionally to remove small variants by filtering out VCF records without “INFO/SVTYPE” values.

Calls were assessed against this benchmark using Truvari^6^ version 4.2.2. The Truvari refine procedure with MAFFT^18^ alignment was used to harmonize query and truth VCF variant representations. The command-line template for Truvari assessment is provided below.

###### Program 4 | Command line template for Truvari assessment against GIAB draft T2T SV benchmark

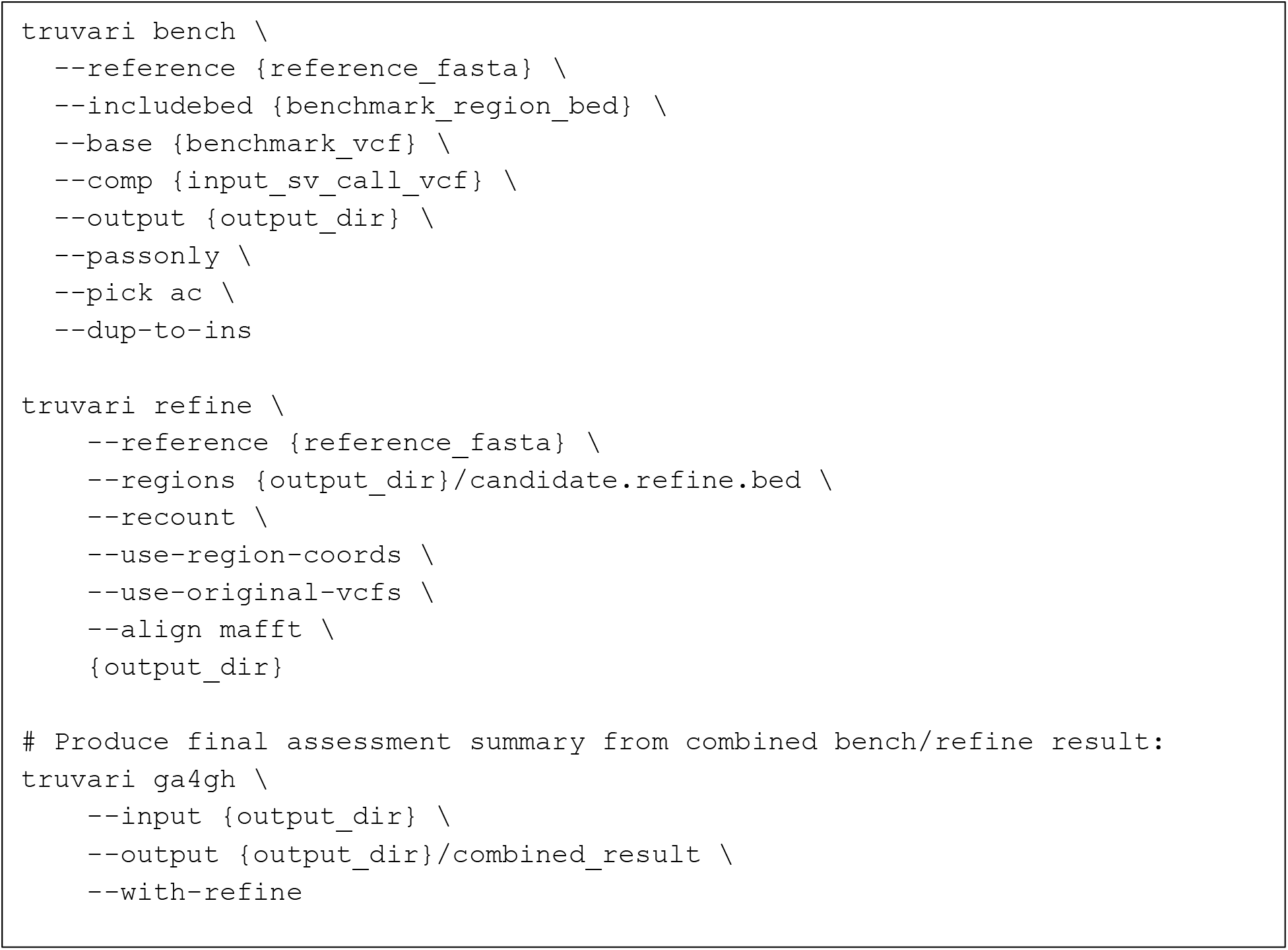

The Truvari refinement procedure described in this command line template relies on phased variant representations from all SV callers. Sawfish already natively provides phased output for very close and overlapping SVs, so to apply this workflow to the additional SV callers we used HiPhase^19^ version 1.4.1 to phase each SV VCF with respect to the sample read alignment file as follows:

###### Program 5 | Command line template for Hiphase-based SV call phasing

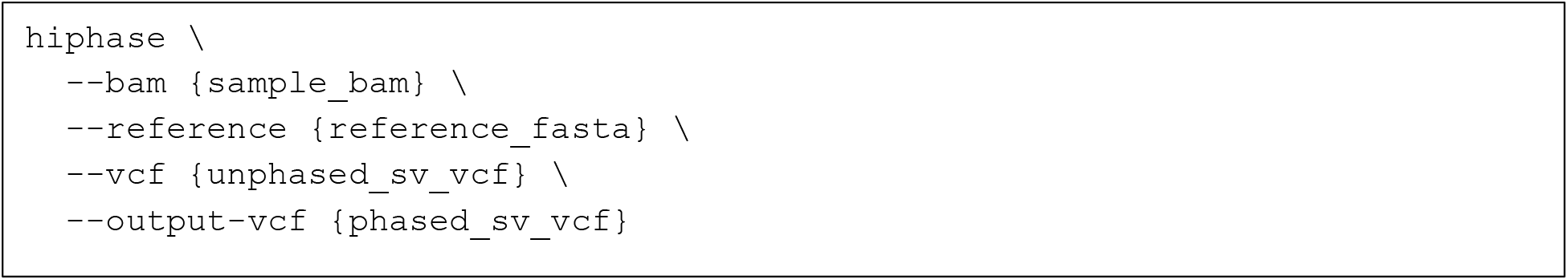

Assessment methods for the GIAB draft T2T SV benchmark are still being developed and optimized. To help check the robustness of our Truvari assessment results, we additionally evaluated all SV calls against this SV benchmark using an additional assessment method, hap-eval^7^, which provides both an independent assessment and uses a technique that does not rely on local SV phasing, such that we can be more confident in the robustness of our SV accuracy observations. For the hap-eval performance described in Results and **Table S1**, we use hap-eval installed from git SHA “e3e2c55” and following the command-line below for each SV caller’s output:

###### Program 6 | Command line template for hap-eval assessment against GIAB draft T2T SV benchmark

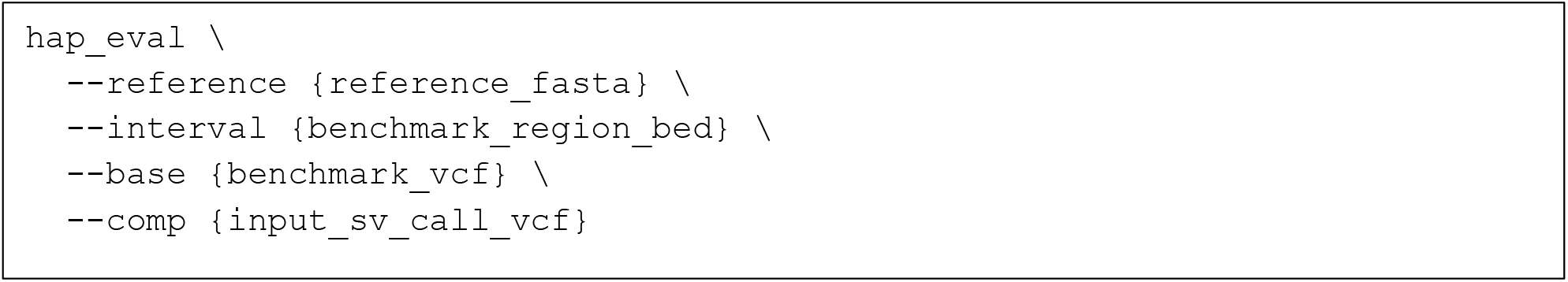

##### Assessment of SV calls against the GIAB CMRG SV benchmark

The GIAB CMRG SV Benchmark 1.0 variant and confidence regions were downloaded from:

https://ftp-trace.ncbi.nlm.nih.gov/ReferenceSamples/giab/release/AshkenazimTrio/HG002_NA24385_son/CMRG_v1.00/GRCh38/StructuralVariant

SV calls were assessed against this benchmark using Truvari version 4.2.2, but with a simplified analysis not requiring the Truvari “refine” method, as described in the command-line template below.

###### Program 7 | Command line template for Truvari assessment against GIAB CMRG SV benchmark

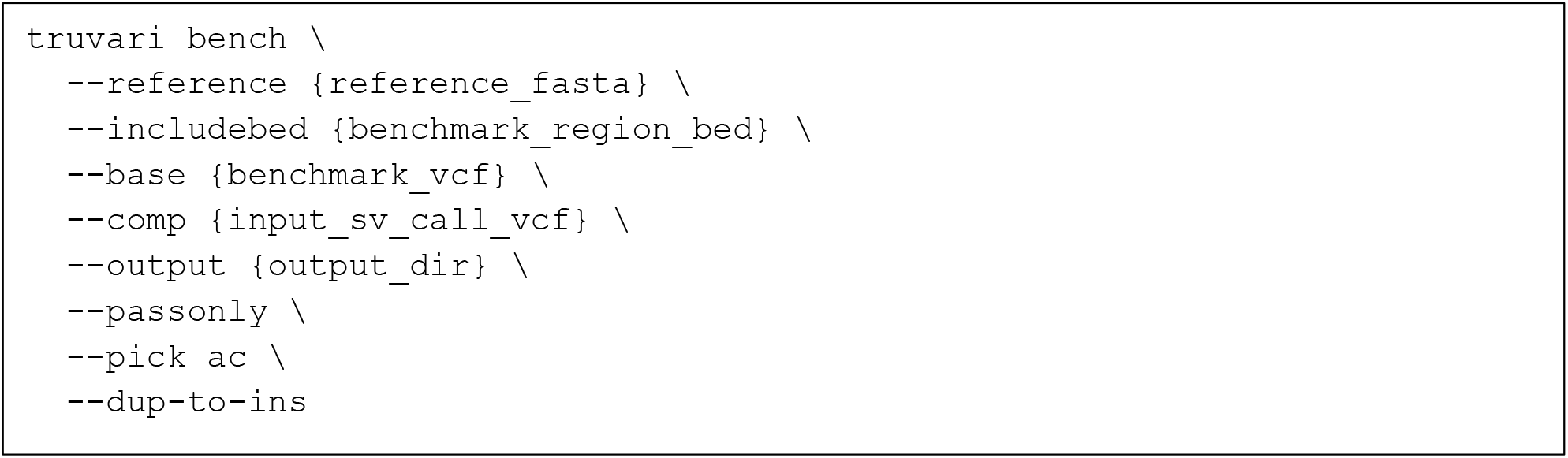

### Joint genotyping accuracy assessment

#### Sequencing Data Processing

We obtained HiFi whole genome sequencing data from all consented samples in CEPH Pedigree 1463 from the recent Porubsky et al. study^14^. All reads were mapped to the GRCh38 reference with pbbm2 version 1.10.0 using the “HIFI” preset option.

#### SV calling

The same SV caller versions used for single-sample calling on HG002 are applied for joint-genotyping on the Platinum Pedigree samples. The command-line options for joint-genotyping on all SV callers are the same as those used for joint-genotyping, with the exception that all samples from CEPH pedigree generations 2 and 3 are included as input to the joint genotyping step using each SV callers sample input format (discovery directory, snf file or svsig file for sawfish, Sniffles2 and pbsv, respectively).

Prior to assessment of joint genotyping concordance, SV joint-genotyping output from all 3 callers are processed in the same way to restrict the evaluated SV set to passing SVs at least 50 bases in length with all breakend records filtered out.

#### Assessment of SV call concordance on the Platinum Pedigree

SV genotype concordance with the Platinum Pedigree is assessed using the pedigree genotype concordance methods described in Porubsky et al.^14^. For each SV call, this concordance evaluation assesses if the genotypes in all samples are concordant with the known pedigree haplotype inheritance pattern for the given region. The “concordance” software used to make this assessment can be compiled and run according to the documentation here:

https://github.com/Platinum-Pedigree-Consortium/Platinum-Pedigree-Inheritance/blob/main/analyses/Concordance.md

Among the evaluated SV callers, we compare both the total count of concordant SV calls and the percentage of all concordant and discordant calls which are assessed as concordant.

### Sawfish SV calling methods

#### Candidate Discovery

The first ‘discover’ step of sawfish is run once on each input sample. In this step each sample is scanned for SV evidence, which are clustered and used to assemble a set of candidate SV alleles. Details of this step are provided below.

##### Scanning sample alignments

The method scans over all mapped reads in the genome to identify clusters of breakpoint-associated read signatures and to create regional depth bins. Reads are filtered from this scan if they are flagged as unmapped, secondary, QC failed or duplicate. Reads with gap compressed identity less than 0.97 are filtered out as well. Reads with mapping quality less than 10 are disqualified from breakpoint evidence scanning but still used to find regional sequencing depth.

Breakpoint evidence is gathered from indel and split read annotations in the read alignments. In each case a simple breakpoint candidate is created with breakends matching the location and orientation implied by the corresponding read alignment feature. Note that split read evidence is parsed from primary alignments only. Additional unpaired breakend evidence is gathered from soft-clipped read edges. Such unpaired breakends are used to assemble large insertions when candidate breakend pairs are found in the expected orientation (details below). Soft-clipped read ends only contribute to breakend candidates when at least 500 bases of the read are soft-clipped on one end, and no clipping is found on the other end of the read.

The average depth of each 2kb bin across the genome is also found while scanning alignments for breakpoint evidence. Gaps created by splitting the read into primary and supplementary alignments are accounted for in the depth calculation, but not the alignment indels.

##### Clustering breakpoint evidence

Breakpoint evidence form individual reads is clustered into candidate breakpoint clusters as follows: Each breakpoint evidence observation is treated as a cluster with a supporting read count of 1. Breakpoint clusters with a matching breakend orientation are tested for their total breakpoint distance, defined as the sum of the distance between each of their breakends. If the breakpoint distance between the clusters is 500 bases or less, the breakpoint clusters are merged, such that each merged breakend extends from the minimum of the two breakend start positions, and the maximum of the two breakend end positions. The merged supporting read evidence of a cluster is the sum of the two input clusters. After merging is completed, clusters with only a single supporting read are discarded as noise. All others comprise the candidate breakpoint cluster set.

##### Candidate breakpoint cluster refinement

In the breakpoint cluster refinement process, candidate breakpoints are assembled into SV haplotype contigs, which are then aligned back to the genome to generate candidate SVs used in downstream merging and genotyping steps.

Refinement begins by defining the regions of the genome used to extract reads for SV contig assembly for each breakpoint candidate. For breakpoint candidates with distant breakends, these will simply be the two breakend regions. For breakpoint candidates that form an indel-like breakend orientation pattern with a breakend distance of 600 bases or less, the assembly region is merged to span the full region between the two breakends. These become single-region candidate breakpoints. Such breakpoints go through an additional clustering step to consolidate all single-region candidate breakpoint regions within 300 bases of each other into a single candidate assembly region, except that the consolidation process is limited to prevent the creation of consolidated regions larger than 8000 bases.

##### Large insertion candidate generation

Large insertions can only be discovered by the standard breakpoint clustering process if the read mapper represents large insertions in the reported read alignments. Additional candidate large insertions can be identified from local soft-clipped read alignment patterns as follows. When a pair of left-anchored and right-anchored soft-clipped breakend candidates are found such that the left and right breakends are within 500 bases, these are converted into large insertion candidate assembly regions if they aren’t already overlapping a candidate assembly region from the standard cluster refinement process.

##### Consensus contig generation

The first step of contig generation is obtaining the subsequences of the reads around each candidate breakpoint position. To do so, reads mapped near the candidate breakpoint are enumerated with the same filtration used for breakpoint evidence discovery. For each read, a trimmed subread is extracted comprising 300 bases of read sequence in each direction from the last base mapped on each side of the candidate breakpoint location. These trimmed reads are additionally filtered out if they lack breakpoint evidence, requiring either an indel at least 25 bases long, or a soft-clipped segment within the trimmed region. Note for the special case of large insertion candidates discovered from paired breakend clusters, no read trimming is applied on the soft-clipped side of the candidate breakend, because the length of the large insertion isn’t known for this case. If there are more than 100 trimmed reads for a given breakpoint candidate, the reads are deterministically subsampled to 100.

The trimmed reads are next clustered and polished into a set of contigs using a simple iterative procedure to assign each read to a partial order alignment (POA) graph using the spoa library^20^. Each of these graphs are generated from previously evaluated trimmed reads, and each is interpreted as being sampled from the same underlying haplotype combined with sequencing noise. Each trimmed read is successively aligned to each POA graph using a linear gap alignment with weights of 1, -3, -1 for match, mismatch, and gap, using wavefront alignment^21^. If at least one read alignment has an aligned read length of 100 or higher, and an alignment score normalized by the aligned read length of 0.96 or higher, then the trimmed read is assigned to the POA graph with the highest normalized alignment score, and the POA graph is updated to include the new trimmed read. If the read does not have a sufficiently high-quality alignment to any existing POA graph, then the read is used to start a new POA graph so long as this would not create more than 8 graphs, otherwise the read is filtered out of the consensus contig generation procedure. After all trimmed reads have been processed, the POA graphs are filtered to remove cases with less than 2 supporting reads, and the top *P* remaining POA graph clusters by supporting read count are used to generate a consensus contig for downstream steps, where *P* is the local ploidy count.

##### Candidate SV Generation

Top contigs generated for each breakpoint cluster are processed for candidate SV information by locally aligning these back to the expected reference locations. For indel-like contigs assembled from reads over a single region of the reference, the contigs can simply be aligned back to a similar segment of the reference sequence and all indels over the minimum size (35 by default) extracted from the alignment as SV candidates.

For all other types of contigs, a synthetic derived chromosome reference is created by appending the two reference region segments corresponding to each candidate breakend location, possibly with one of the reference segments reverse-complemented according to the candidate breakend orientation pattern. In this way the contig can be aligned to the synthetic reference with an apparent large deletion extending between the two reference regions, and this alignment can be processed back into a pair of breakends of any orientation, which generalizes to handle all types of SV breakpoints.

Whether using the small indel or generalized breakpoint reference alignment procedure, each breakpoint alignment can be standardized by left-shifting its position and finding the full breakend homology range and insertion sequence.

#### Joint Calling

The second ‘joint-call’ step of the sawfish pipeline enables candidate SVs to be analyzed across multiple samples. The primary steps are: (1) consolidation of SV candidates which are duplicated across samples; (2) evaluation of read support for the deduplicated candidate SV haplotypes to genotype the SV in each sample; (3) evaluation of depth support for large copy-changing events; and (4) reporting all SVs as a VCF jointly genotyped across all samples.

##### Duplicate Haplotype Consolidation

As a first step to duplicate haplotype merging, overlapping haplotypes from all samples are pooled into groups from which duplicates are found and consolidated. This pooling procedure is run separately within the set of indel-like SV candidates consolidated to a single reference region (as described in the candidate discovery section above), and all other candidates associated with multiple reference regions. For indel-like candidates, the candidate pool is found from all intersecting candidate regions. For all other SV candidates associated with two reference regions, pools are composed of candidates where both reference regions are within 100 bases of at least one other candidate in the pool.

Within each candidate haplotype pool, candidates are clustered into duplicate groups based on pairwise testing first for matching breakpoints, then for very high haplotype sequence similarity if the breakpoints aren’t an exact match, with an exception made to exclude duplicate merging of haplotypes candidates from the same sample. Haplotype sequence similarity is determined by using a linear gap aligner with 1, -3, -2 for match, mismatch, and gap scores. If the resulting alignment score normalized by the aligned haplotype length is at least 0.97 then the haplotypes are treated as duplicates. Within each duplicate haplotype group, one member is chosen as representative based on having the highest supporting read count from contig assembly, or longest contig size when supporting read count is tied.

##### Genotyping

SVs are genotyped in the context of the overlapping haplotype pools created for the purpose of duplicate haplotype identification. Each breakend of each SV allele is evaluated by aligning segments of locally mapped reads to the corresponding segment of the SV haplotype assembly, in addition to aligning these reads to the reference sequence and other SV alleles in the overlapping haplotype pool. The segments of the read, reference and SV haplotypes selected for this purpose on each candidate SV breakend are extended up to 500 bases from the non-anchored edge of the breakend’s homology range. Alignment is scored using a linear gap aligner with 1, - 3, -2 for match, mismatch, and gap scores, with alignment scores normalized by the aligned read segment length.

The alignment scores for each read are next converted into support counts, where each read could be identified as uniquely consistent with either the haplotype of the SV allele in question, the reference haplotype or a candidate overlapping SV haplotype. Candidate overlapping haplotypes are only identified for local indel-like SV candidates. The overlapping case can be from a second SV haplotype assembled from reads in the given sample. If a second haplotype wasn’t assembled for the sample, then overlapping SV haplotypes from other samples (among those remaining after merging duplicate haplotypes) are compared to find the overlapping SV haplotype with the highest support from all sample reads – if such an overlapping case is found it is added as a second ‘guest’ candidate haplotype.

Each read’s support for the SV haplotype is evaluated separately for each breakend of a given breakpoint. A read is counted as supporting an SV allele if its alignment score to the SV haplotype is better than the reference haplotype for any breakend. A read can only support the reference haplotype if the reference haplotype alignment has a higher score on all evaluated breakends. This arrangement helps to counteract various forms of reference bias in the alignment scores. Reads supporting an overlapping SV haplotype are internally recorded for the overlapping haplotype but converted into reference allele support for both quality score calculations and in the final VCF allele count output, per the reporting convention used by most available SV calling tools.

##### Quality model

The sawfish quality scores are generated from a simplified model which generates qualities directly from the read support counts for each allele. Ideally the scores of read alignments to each allele haplotype would be used instead of counts, but thus far the count approximation hasn’t been attributed to any substantial fraction of SV genotyping errors.

For each SV allele and each sample, we solve for the posterior probability of diploid genotypes, *G*, given the observed read counts at the SV loci, *D* as follows

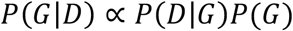

The diploid genotypes are *G* = {*ref, het, hom*} representing 0,1 or 2 copies of the SV allele. The genotype prior is

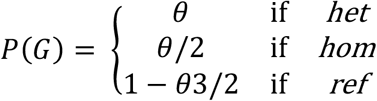

where *θ* = 5 x 10^-4^.

The genotype likelihood is found as follows

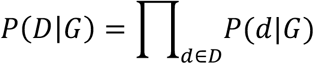

treating each read observation *d* ∈ *D* as independent. The read likelihood is

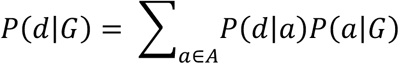

where *A* = {*ref, alt*} are the SV alleles in the model representing the reference and SV haplotypes. As previously discussed, support for any overlapping SV haplotypes is counted towards the reference allele for the purpose of the genotype quality model, which is intentionally simplified to represent only one SV haplotype at a time.

Considering the terms in the read likelihood, the allele likelihood *P(d*|*a)* is set from the read allele support counts using a single erroneous read support count probability *e* = 1 x 10^-5^ for all cases. The allele probabilities *P(a*|*G)* are the simple allele fractions (0, 0.5, 1) associated with each genotype.

##### Evaluating read coverage support for larger SV calls

After the above described read-based genotyping steps, all deletion and duplication candidates of at least 50kb are additionally evaluated for an SV depth signature consistent with the SV type. If the supporting SV depth signature is not found, the corresponding SV breakpoint is still reported in the VCF output, but as a set of breakend (BND) records, rather than as a deletion or duplication. In this way, breakpoints comprising larger multi-breakpoint complex SVs can still be reported in the VCF output without compromising the precision of simpler copy-number changing deletion and tandem duplication events.

The criteria for a consistent depth signature in each sample is based on the average depth of the interior region of the SV is compared to the average depth in the 6kb regions to the left and right flank. The ratio of the interior to flank depth must be no more than 0.8 for a deletion and no less than 1.2 for a duplication. In a multi-sample context, a consistent depth signature is required in at least half of samples with a non-reference genotype to retain the SV type as a deletion or duplication. While this approach applies a very simple heuristic to assess SV support for each large SV, this has been found to be effective in filtering typical complex SV breakpoints from true large deletion and duplication signatures.

## Supplementary Figures

**Figure S1.**
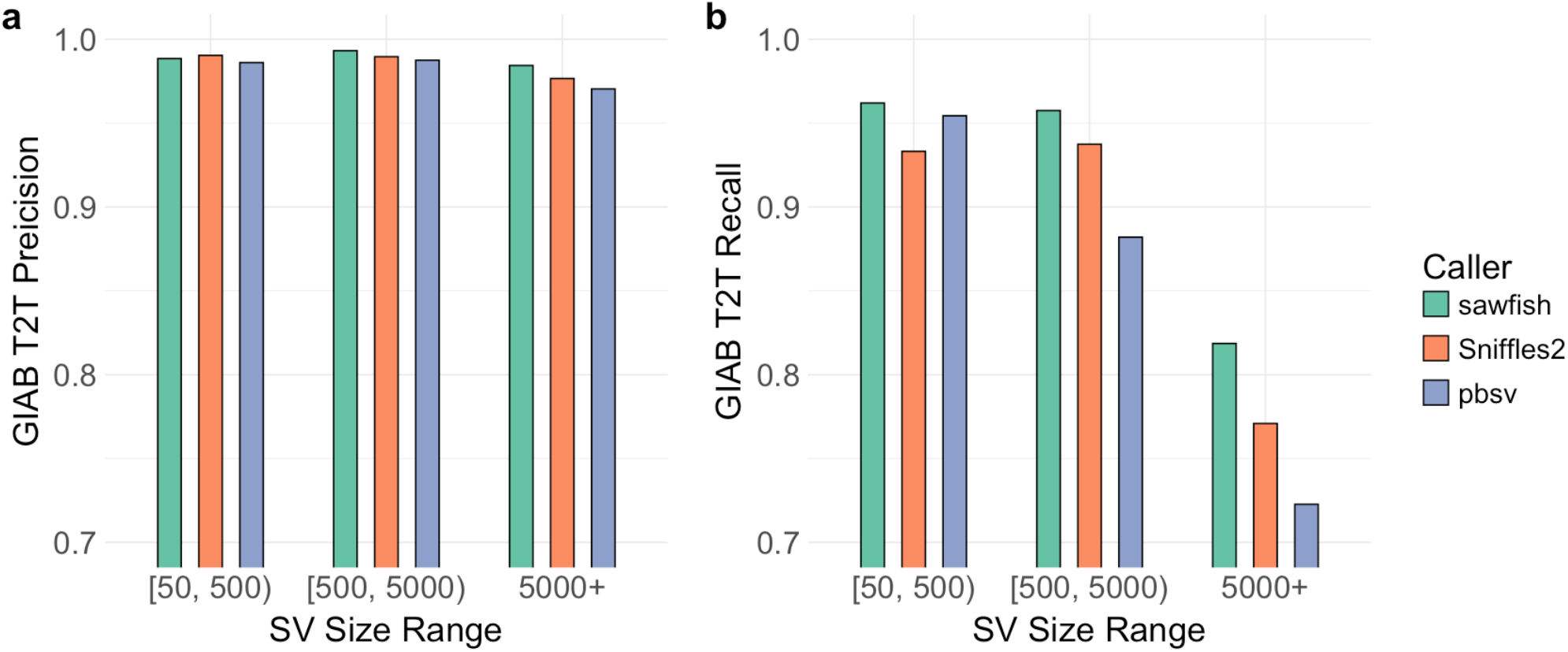
SV caller accuracy assessment as a function of SV size. Assessment of all SV callers against the GIAB T2T benchmark set stratified into 3 SV size categories. Benchmarking details are as described for **Figure 1a** but showing precision (a) and recall (b) instead of F1-score.

**Figure S2.**
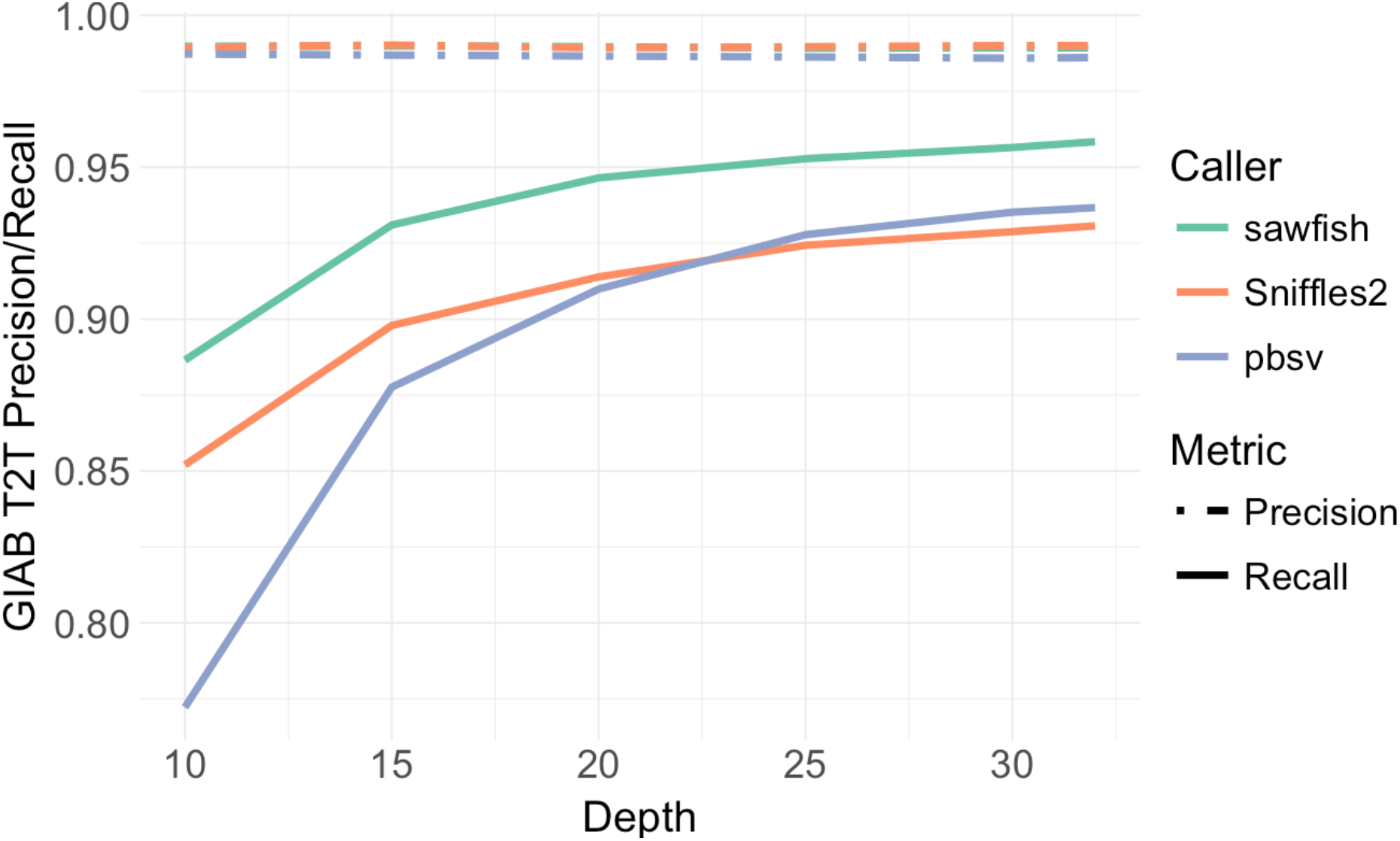
SV caller accuracy assessment as a function of depth. Assessment of all SV callers against the GIAB T2T benchmark set for aligned read inputs subsampled from 32-fold to 10-fold coverage. Benchmarking details are as described for **Figure 1b** but showing precision and recall values instead of F1-score.

## Supplementary Tables

**Table S1.** SV assessment results on GIAB draft T2T SV benchmark using Truvari or hap-eval for 32-fold (full) WGS coverage input.

| SV Caller | Assessment Method | F1-Score | Recall | Precision |
| --- | --- | --- | --- | --- |
| sawfish | Truvari | 0.9736 | 0.9584 | 0.9893 |
| Sniffles2 | Truvari | 0.9595 | 0.9307 | 0.9900 |
| pbsv | Truvari | 0.9608 | 0.9367 | 0.9861 |
| sawfish | hap-eval | 0.9739 | 0.9658 | 0.9822 |
| Sniffles2 | hap-eval | 0.9387 | 0.9166 | 0.9620 |
| pbsv | hap-eval | 0.9323 | 0.9240 | 0.9408 |

**Table S2.**
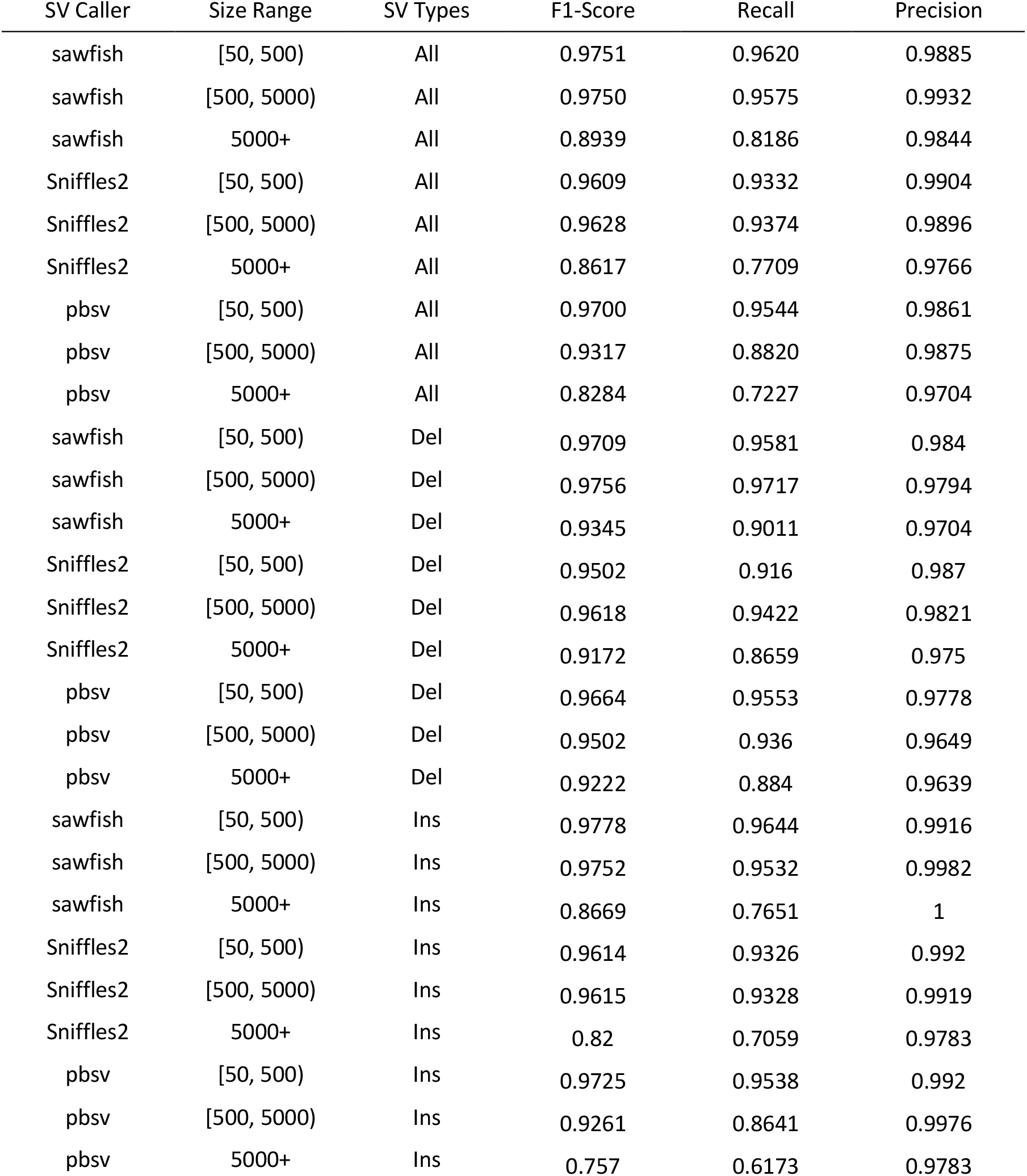
SV assessment results on GIAB draft T2T SV benchmark using Truvari with results binned into 3 SV size ranges, and 3 SV type groups for all SVs (All), deletions (Del), or insertions (Ins).

| SV Caller | Size Range | SV Types | F1-Score | Recall | Precision |
| --- | --- | --- | --- | --- | --- |
| sawfish | [50, 500) | All | 0.9751 | 0.9620 | 0.9885 |
| sawfish | [500, 5000) | All | 0.9750 | 0.9575 | 0.9932 |
| sawfish | 5000+ | All | 0.8939 | 0.8186 | 0.9844 |
| Sniffles2 | [50, 500) | All | 0.9609 | 0.9332 | 0.9904 |
| Sniffles2 | [500, 5000) | All | 0.9628 | 0.9374 | 0.9896 |
| Sniffles2 | 5000+ | All | 0.8617 | 0.7709 | 0.9766 |
| pbsv | [50, 500) | All | 0.9700 | 0.9544 | 0.9861 |
| pbsv | [500, 5000) | All | 0.9317 | 0.8820 | 0.9875 |
| pbsv | 5000+ | All | 0.8284 | 0.7227 | 0.9704 |
| sawfish | [50, 500) | Del | 0.9709 | 0.9581 | 0.984 |
| sawfish | [500, 5000) | Del | 0.9756 | 0.9717 | 0.9794 |
| sawfish | 5000+ | Del | 0.9345 | 0.9011 | 0.9704 |
| Sniffles2 | [50, 500) | Del | 0.9502 | 0.916 | 0.987 |
| Sniffles2 | [500, 5000) | Del | 0.9618 | 0.9422 | 0.9821 |
| Sniffles2 | 5000+ | Del | 0.9172 | 0.8659 | 0.975 |
| pbsv | [50, 500) | Del | 0.9664 | 0.9553 | 0.9778 |
| pbsv | [500, 5000) | Del | 0.9502 | 0.936 | 0.9649 |
| pbsv | 5000+ | Del | 0.9222 | 0.884 | 0.9639 |
| sawfish | [50, 500) | Ins | 0.9778 | 0.9644 | 0.9916 |
| sawfish | [500, 5000) | Ins | 0.9752 | 0.9532 | 0.9982 |
| sawfish | 5000+ | Ins | 0.8669 | 0.7651 | 1 |
| Sniffles2 | [50, 500) | Ins | 0.9614 | 0.9326 | 0.992 |
| Sniffles2 | [500, 5000) | Ins | 0.9615 | 0.9328 | 0.9919 |
| Sniffles2 | 5000+ | Ins | 0.82 | 0.7059 | 0.9783 |
| pbsv | [50, 500) | Ins | 0.9725 | 0.9538 | 0.992 |
| pbsv | [500, 5000) | Ins | 0.9261 | 0.8641 | 0.9976 |
| pbsv | 5000+ | Ins | 0.757 | 0.6173 | 0.9783 |

**Table S3.** SV assessment results on GIAB draft T2T SV benchmark using Truvari for 32-fold (full) WGS coverage and subsampled levels down to 10-fold coverage.

| SV Caller | Coverage | F1-Score | Recall | Precision |
| --- | --- | --- | --- | --- |
| sawfish | 10 | 0.9353 | 0.8865 | 0.9898 |
| sawfish | 15 | 0.9595 | 0.9310 | 0.9899 |
| sawfish | 20 | 0.9676 | 0.9465 | 0.9897 |
| sawfish | 25 | 0.9707 | 0.9528 | 0.9892 |
| sawfish | 30 | 0.9727 | 0.9565 | 0.9893 |
| sawfish | 32 | 0.9736 | 0.9584 | 0.9893 |
| Sniffles2 | 10 | 0.9156 | 0.8520 | 0.9895 |
| Sniffles2 | 15 | 0.9418 | 0.8979 | 0.9902 |
| Sniffles2 | 20 | 0.9501 | 0.9139 | 0.9893 |
| Sniffles2 | 25 | 0.9559 | 0.9243 | 0.9897 |
| Sniffles2 | 30 | 0.9584 | 0.9288 | 0.9900 |
| Sniffles2 | 32 | 0.9595 | 0.9307 | 0.9900 |
| pbsv | 10 | 0.8666 | 0.7722 | 0.9873 |
| pbsv | 15 | 0.9291 | 0.8776 | 0.9869 |
| pbsv | 20 | 0.9467 | 0.9099 | 0.9866 |
| pbsv | 25 | 0.9562 | 0.9278 | 0.9863 |
| pbsv | 30 | 0.9599 | 0.9352 | 0.9859 |
| pbsv | 32 | 0.9608 | 0.9367 | 0.9861 |

**Table S4.** SV assessment results on GIAB Challenging Medically Relevant Genes (CMRG) benchmark. Assessment results of each SV caller against the GIAB CMRG benchmark using Truvari. FN: False Negative, FP: False Positive.

| SV Caller | F1-Score | Recall | Total<br>FNs | Precision | Total FPs |
| --- | --- | --- | --- | --- | --- |
| sawfish | 0.9929 | 0.9907 | 2 | 0.9951 | 1 |
| Sniffles2 | 0.9640 | 0.9630 | 8 | 0.9650 | 7 |
| pbsv | 0.9714 | 0.9583 | 9 | 0.9849 | 3 |

**Table S5.** Pedigree concordant SV calls on CEPH-14633 generation 2 and 3.

| SV Caller | Total SVs | Concordant SVs | Discordant SVs | % Concordant |
| --- | --- | --- | --- | --- |
| sawfish | 41026 | 31956 | 9070 | 77.89 |
| Sniffles2 | 33034 | 21548 | 11486 | 65.23 |
| pbsv | 30769 | 22841 | 7928 | 74.23 |
| sawfish HQ (GQ >= 40) | 32326 | 27877 | 4449 | 86.24 |
| Sniffles2 HQ (GQ >= 10) | 8965 | 7643 | 1322 | 85.25 |

